# Quantifying Uncertainty in Phasor-Based Time-Domain Fluorescence Lifetime Imaging Microscopy

**DOI:** 10.1101/2025.04.04.647302

**Authors:** Qinyi Chen, Jongchan Park, Shuqi Mu, Liang Gao

## Abstract

The phasor approach to time-domain fluorescence lifetime imaging microscopy (FLIM) offers a powerful, fit-free method for analyzing complex fluorescence decay signals. However, its quantitative accuracy is fundamentally limited by noise—particularly photon shot noise—which introduces variability and bias in lifetime estimation and fluorophore unmixing. In this study, we present a theoretical uncertainty model for phasor-based time-domain FLIM that analytically captures the propagation of shot noise and quantifies its impact on phasor coordinates and fluorophore weight estimation. We validate the model using Monte Carlo simulations and experimental data acquired from standard fluorescent dyes and biological tissue samples. Our model improves the overall reliability and efficiency of phasor-based time-domain FLIM, particularly in photon-limited imaging applications.

## 1. Introduction

The phasor approach has emerged as a powerful and fit-free method for analyzing time-domain Fluorescence Lifetime Imaging Microscopy (FLIM) data[1,2]. Unlike conventional fitting-based methods, it transforms fluorescence decay signals into a two-dimensional phasor space using a normalized Fourier transform[3]. This transformation streamlines lifetime analysis by providing a fast, intuitive graphical representation. Each decay curve is mapped to a point on the phasor plot with coordinates (G, S), enabling straightforward discrimination of fluorophores with distinct lifetimes—without requiring assumptions about the underlying decay kinetics[4].

One of the key advantages of the phasor method is its ability to separate multiple fluorescent species within a single pixel, leveraging the principle of phasor addition[5]. In this framework, the phasor of a mixed fluorescence signal lies on the linear trajectory connecting the phasors of the individual species, with its position reflecting their relative contributions[6]. This linearity enables robust and model-independent lifetime unmixing. As a result, the phasor approach has found widespread applications in metabolic imaging[7], Förster resonance energy transfer (FRET) measurements[8], and studies of protein-protein interactions[9], where quantitative lifetime mapping provides valuable biological insights.

Previous research on phasor plot analysis has predominantly focused on the segmentation, clustering, and unmixing of point clouds using methods such as principal component analysis (PCA)[10,11], k-means clustering[12], Gaussian mixture model (GMM)-based soft clustering[13], and deep learning approaches[14–16]. Analyses of point cloud shape has primarily relied on multi-parameter assessment[17] or explored its inverse correlation with photon count[18]. Despite these advances, the FLIM phasor approach remains highly susceptible to uncertainties introduced by experimental noise and photon counting limitations[19]. Critically, there is still no comprehensive frameworks that systematically characterize the influence of noise on phasor-based lifetime measurements or model the statistical behavior of point clouds in the phasor space.

Understanding the sources and propagation of uncertainty in time-domain FLIM phasor analysis is essential for improving the reliability of lifetime unmixing and their biological interpretations. Among available time-domain FLIM techniques[20], time-correlated single photon counting (TCSPC) provides the highest temporal resolution for resolving complex, multi-exponential fluorescence decays[21]. This fine resolution is achieved by recording the arrival time of the first detected photon in each excitation cycle, enabling the construction of a temporal histogram with a large number of narrow time bins[22]. However, such discretization reduces the number of detected photons per bin[23], making the measurements particularly vulnerable to shot noise — the statistical uncertainty inherent to the discrete and stochastic nature of photon detection. Consequently, shot noise becomes the dominant source of error in phasor coordinate estimation, leading to unavoidable statistical fluctuations. While increasing the total photon count can improve the signal-to-noise ratio (SNR), the fundamental stochasticity of photon arrival times means that shot-noise–induced errors in the phasor plot can never be neglected.

In this study, we present a quantitative framework for evaluating uncertainty in time-domain FLIM phasor analysis by developing a theoretical model that captures noise propagation and measurement variability. We validate this model using both simulations and experimental data from well-characterized fluorescent dyes, and further demonstrate its applicability to real biological specimens. By systematically evaluating the effects of noise and acquisition parameters on phasor-based lifetime estimation, our work provides a foundation for improving the accuracy, reliability, and interpretability of time-domain FLIM in biological imaging.

## 2. Method

### 2.1 Uncertainty Model

TCSPC-based FLIM[24] records only the first detected photon per excitation cycle, enabling accurate photon counting under low-light conditions. Both excitation and subsequent photon emission are independent stochastic processes, resulting in probabilistic photon arrival times. By repeatedly exciting the sample over multiple cycles, TCSPC accumulates photon arrival events into a temporal histogram distributed across discrete time bins. Since each photon detection event is independent and the probability of detection per cycle is low, the number of photons detected within each time bin follows a Poisson distribution, where the variance equals the mean[25]. This intrinsic statistical property propagates Poisson noise into the phasor space, ultimately impacting the precision of fluorescence lifetime estimation and the decomposition of fluorophore components.

For an ideal FLIM measurement without noise, the phasor coordinates (*G,S*) at a given pixel are computed by applying a normalized Fourier transform to the fluorescence decay histogram:

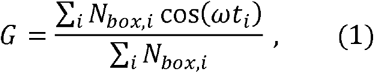

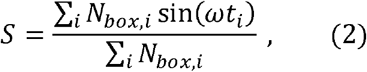

where *N*_*box,i*_ represents the photon count detected at time bin *t*_*i*_, and *ω* is the angular frequency corresponding to the analysis temporal histogram range, usually set to the first harmonic. However, due to shot noise, the observed photon count *N*_*noisy,i*_ in each time bin follows a Poisson distribution:

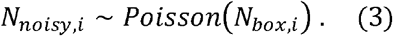

A defining property of the Poisson distribution is that its variance equals its mean:

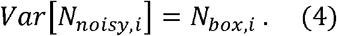

Shot noise causes phasor coordinates to fluctuate randomly, leading to systematic deviations in lifetime estimation and fluorophore unmixing. To quantify these deviations, we apply error propagation theory to the phasor transformation equations (**Supplementary 1**). The variance and covariance of the noisy phasor coordinates can be analytically expressed as:

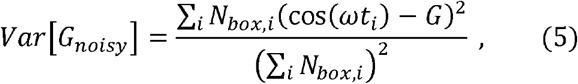

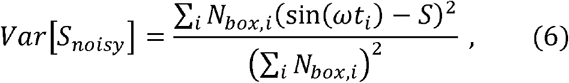

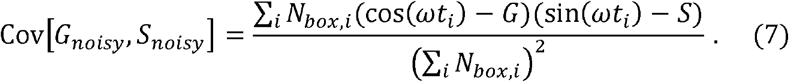

These expressions imply that, even under ideal single-component decay conditions, phasor coordinates exhibit a two-dimensional Gaussian distribution centered at the noise-free value. In the case of multi-component mixtures, where the phasor of each species contributes linearly to the overall signal, shot noise can shift the measured phasor away from its expected position, introducing uncertainty into component weight estimation.

To model this effect, we assume the phasor coordinates follow a multivariate normal distribution. The probability density function for observing a noisy phasor coordinate ***x*** near the expected mean *μi* is given by:

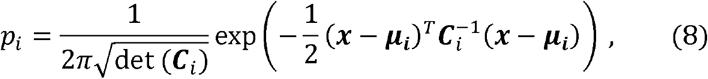

where *C*_*i*_ is the covariance matrix of the distribution, representing shot-noise-induced uncertainty, and *μ*_*i*_ is the expected phasor location for fluorophore *i*. By normalizing these likelihoods across candidate components, we compute the probability that the observed phasor corresponds to each fluorophore:

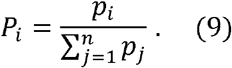

Based on this theory, we developed an uncertainty-aware phasor analysis workflow consisting of the following steps:

1. Compute total photon counts and phasor coordinates for each pixel based on fluorescence decay histograms.
2. Estimate probable fluorophore fractions using the linear combination principle in the phasor plot.
3. Determine the uncertainty range by incorporating shot noise-induced variance from the Poisson model.
4. Apply a Gaussian probability model to assign statistical confidence to each estimated fluorophore fraction.
5. Refine mixture weight estimation by normalizing probabilities across multiple potential fluorophore distributions.

This workflow allows for a robust estimation of fluorescence lifetimes and mixture compositions by accounting for noise-induced variability. By integrating probabilistic models into phasor-based FLIM analysis, we can improve the reliability of results, particularly in low-photon or mixed-lifetime imaging scenarios.

### 2.2 Materials

To validate the proposed uncertainty model, two standard fluorescent dyes were utilized: Rhodamine 110 (98+%) and Alexa Fluor™ Carboxylic Acid, tris(triethylammonium) salts (ThermoFisher)[26]. The dyes were dissolved in phosphate-buffered saline (PBS, pH 7.2, ThermoFisher) at a low concentration of approximately 1 × 10^−4^ *μg/μl*. The prepared solutions were mixed in predetermined ratios to 100*μl* a final volume of and subsequently loaded into well plates for imaging.

Additionally, the uncertainty evaluation model was applied to a biological sample—a prepared mouse kidney tissue section (FluoCells Prepared Slide, ThermoFisher). This sample was stained with Alexa Fluor 488 wheat germ agglutinin (WGA) to label cell membranes and Alexa Fluor 568 phalloidin to stain filamentous actin (F-actin). These fluorophores exhibit distinct but closely spaced fluorescence lifetimes (Alexa Fluor 488: 2.27 ns; Alexa Fluor 568: 2.69 ns), making them suitable for validating the model’s sensitivity to lifetime differences. Application of the uncertainty model enabled a quantitative assessment of fluorophore fraction estimation and associated confidence levels within the biological tissue.

### 2.3 Imaging Setup

Fluorescence lifetime imaging was performed using a Leica SP8 FLIM confocal microscope. For imaging the two fluorescent dyes, excitation was provided by an 80LMHz pulsed laser source at 497□nm (for Rhodamine 110) and 647□nm (for Alexa Fluor™ Carboxylic Acid, tris(triethylammonium) salts), with fluorescence emission collected in the spectral ranges of 500–600□nm and 650–700□nm, respectively. For the mouse kidney tissue section, pulsed excitation lasers at 488□nm and 580□nm were used to excite Alexa Fluor 488 and Alexa Fluor 568, respectively. Emitted fluorescence was collected in the corresponding detection bands of 500–570□nm (Alexa Fluor 488) and 600–651□nm (Alexa Fluor 568). Photon arrival times were recorded across the relevant detection channels using time-correlated single photon counting (TCSPC), enabling the construction of fluorescence decay histograms for subsequent phasor-based FLIM analysis.

To systematically investigate the effect of total photon count on fluorophore weight uncertainty in the mouse kidney section, we varied the number of frame repetition (i.e., summation) rather than adjusting laser power or detector gain. This approach avoids introducing photobleaching to the sample or altering the system’s noise characteristics. Increasing frame repetition effectively extends the total exposure time, allowing for controlled photon accumulation while maintaining consistent imaging conditions.

## 3. Results

### 3.1 Simulations

We validate our analytical uncertainty model by comparing it with Monte Carlo simulations at three distinct photon count levels: low (500 photons), medium (1000 photons), and high (2000 photons). For the two-component case, the fluorescence lifetimes of the pure components were set to 1 ns and 4 ns, with photons mixed in a 50:50 ratio. For the three-component case, the fluorescence lifetimes were set to 0.3 ns, 1 ns, and 4 ns, with a respective ratio of 20:30:50. In both cases, the acquisition frequency was fixed at 200 MHz to ensure consistency in data collection and facilitate comparison of uncertainty across different photon count levels.

Based on the properties of the phasor plot, the fluorescence decay histogram of a single pixel maps to a single point in the phasor plot. In our simulation, we first generated ideal fluorescence decay histograms under the above conditions, representing the noise-free case. To simulate the effect of shot noise, we applied Monte Carlo sampling by drawing photon counts for each time bin from a Poisson distribution with a mean equal to the corresponding noise-free value. This process was repeated for 10,000 iterations to obtain a statistically robust phasor point cloud distribution.

The resulting two-dimensional histograms are shown in Fig 2 (A1-A3, C1-C3), visualizing the spread of phasor points due to shot noise. The red ellipse represents the three-standard-deviation confidence interval predicted by the uncertainty model. As shown in Fig 2 (B1, B2, D1, D2), the bar plots depict the count of simulated points falling within each interval, while the overlaid curves represent the corresponding probability distributions derived from the uncertainty model. The close agreement between simulated data and theoretical predictions confirms that the uncertainty model captures the statistical variability introduced by shot noise.

**Fig. 1.**
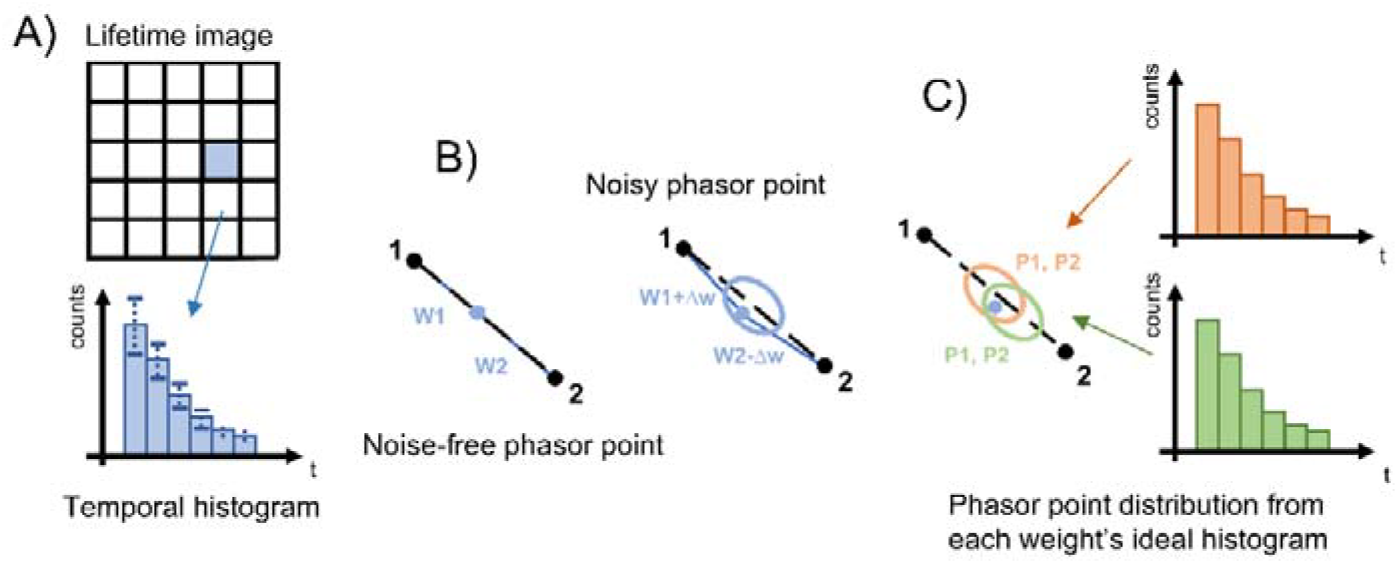
Phasor-based uncertainty analysis in time-domain FLIM. (A) Each pixel in the FLIM image corresponds to a temporal fluorescence decay histogram, which is mapped to a specific point in the phasor plot. (B) Due to the presence of shot noise, the observed phasor point deviates from its noise-free counterpart. This deviation follows a two-dimensional normal distribution centered around the noise-free point. (C) The uncertainty of a noisy phasor point is estimated by assessing its likelihood of belonging to multiple normal distributions, each representing different pair of component weights.

**Fig. 2.**
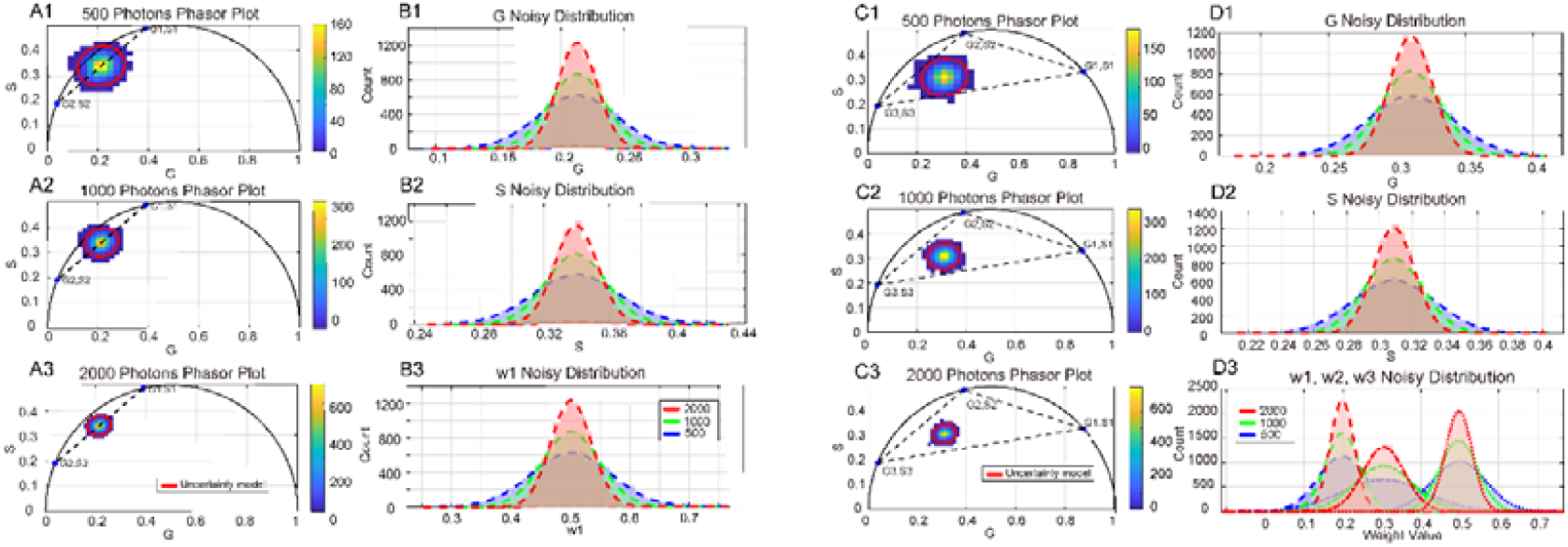
Monte Carlo simulation and uncertainty model’s three-standard-deviation confidence interval prediction based on shot noise in three different total photon levels. (A1-A3, C1-C3) Two-component and three-component point cloud from Monte Carlo simulation and the three-standard-deviation confidence interval predicted by the uncertainty model in the phasor plot. (B1, B2, D1, D2) G and S distributions from Monte Carlo simulation (histogram) and uncertainty model (curve). (B3, D3) Weight distribution for pure components from Monte Carlo simulation.

As illustrated in the weight distribution curve in Fig 2 (B3, D3), increasing the total photon count reduces the impact of shot noise on the displacement of phasor points. Consequently, the estimated component weight distributions converge more closely to the theoretical mixing ratios of 50:50 and 20:30:50.

### 3.2 Experiments

To experimentally validate the uncertainty model, we first analyzed the phasor distribution of a single dye under three different excitation light intensities. As shown in Fig 3 (A1-3, B1, B2), the experimentally measured (G, S) coordinates from 400 pixels are visualized using a blue-to-yellow color gradient, while the red ellipses denote the predicted three-standard-deviation confidence intervals from the uncertainty model. As laser intensity increases, the spread of the phasor coordinates progressively narrows, reflecting improved SNR. The close agreement between the measured data and the model predictions across all conditions confirms the model’s accuracy in quantifying shot-noise-induced variability.

**Fig. 3.**
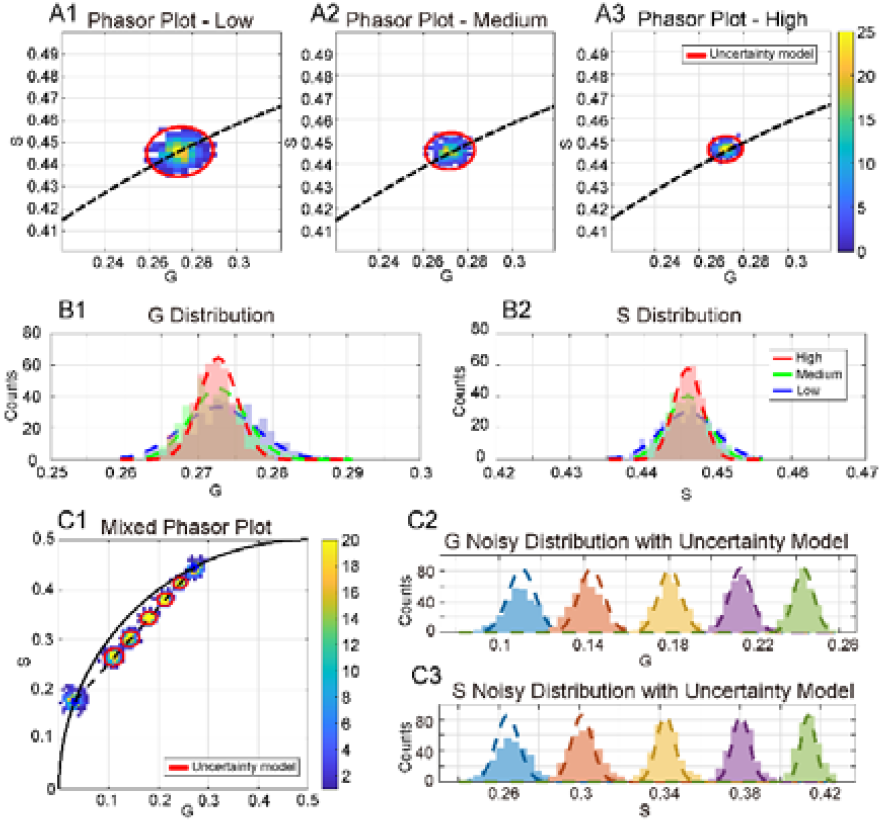
Validating the uncertainty model using fluorescent dyes. (A1-A3) Single dye’s experimental point cloud and three-standard-deviation confidence interval from the uncertainty model at three different laser power levels. (B1, B2) G and S distributions of the single-dye phasor data: experimental (bar histogram) vs. uncertainty model (curve). (C1) Two dyes’ experimental point cloud and three-standard-deviation confidence interval from the uncertainty model at five different mixing ratios. (C2, C3) G and S distributions of the two-dye phasor data: experimental (bar histogram) vs. uncertainty model (curve).

We then examined the phasor distribution of two mixed dyes at five different mixing ratios. As illustrated in Fig 3 (C1-C3), the experimentally measured (G, S) distributions and the corresponding predictions from the uncertainty model are visualized using the same color scheme and confidence ellipses as previously described.

Ideally, the three-standard-deviation confidence ellipses should enclose 99% of the data points; however, in practice, they contain only about 92%. This discrepancy may be attributed to additional experimental noise sources—such as electronic noise, instrument instability, and environmental fluctuations— that are not explicitly modeled. While these factors may contribute to the observed spread, they are not considered the sources of error and are thus not incorporated into the current uncertainty framework.

Next, we applied our uncertainty model to quantify the weight uncertainty of two pure components that exhibit close fluorescence lifetimes in a biological mouse kidney tissue section. Imaging was performed at three different total photon levels to assess how photon statistics influences the uncertainty in component estimation. To investigate the spatial distribution of the components, two regions of interest (ROIs) were selected for analysis.

In the first ROI (Fig 4 A1-A3), the two lifetime components are fully intermixed, with most pixels containing contributions from both species. In contrast, the second ROI (Fig 4 A4-A6) displays a more spatially segregated distribution, where most pixels are dominated by photons from a single fluorophore. These panels display pseudo-colored lifetime maps, where brightness represents fluorescence intensity and hue encodes lifetime values.

**Fig 4.**
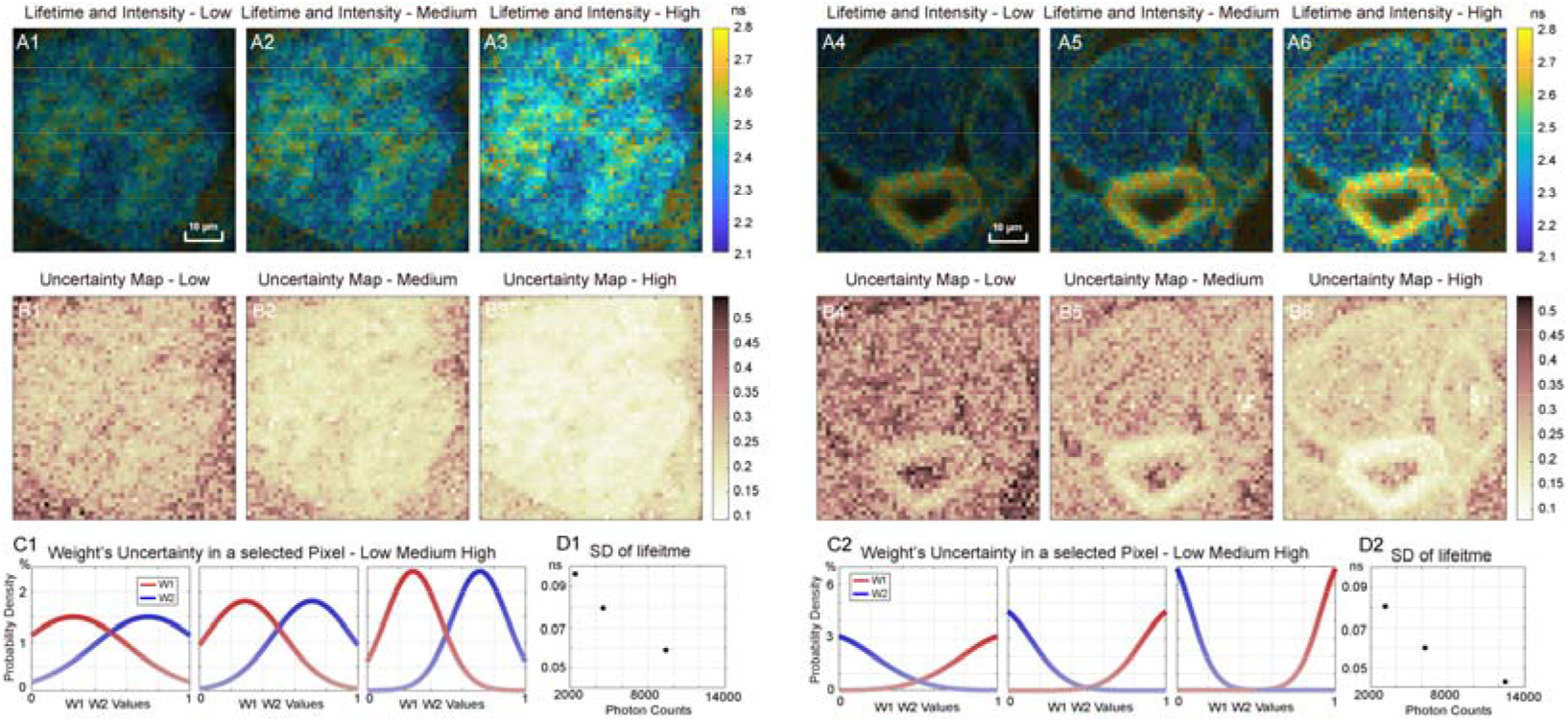
Weight uncertainty analysis of a mouse kidney section. (A1-A6) Lifetime images of ROI 1 and ROI 2 overlaid with intensity images at three photon levels. (B1-B6) Standard deviation maps of weight uncertainty for the two regions. (C1, C2) Weight probability distributions for two fluorophore components for a representative pixel. (D1, D2) Standard deviations of lifetimes at varied total photon counts.

For the uncertainty calculation, we defined 100 equally spaced weight values ranging from 0 to 1 and computed their probability distribution. As shown in Fig. 4 (A1–A6), the intensity and lifetime maps for three photon count levels—with relative ratios of 1:2:4—are presented for both ROIs. The corresponding weight uncertainty maps of fluorophore Alexa Fluor 488, derived from our model, are illustrated in Fig. 4 (B1–B6). In these maps, the color bar indicates the standard deviation of the estimated fluorophore weight for each pixel, providing a quantitative measure of uncertainty. Higher standard deviation values correspond to greater ambiguity in component estimation due to limited photon statistics.

At the pixel level, Fig 4 (C1, C2) shows the uncertainty curves of the weight distribution for both fluorophore components, while Fig 4 (D1, D2) depicts the standard deviation of fluorescence lifetime for a representative pixel. As expected, increasing photon counts enhances the accuracy of both weight and lifetime estimations. Our uncertainty model therefore provides a practical framework for optimizing experimental parameters—such as laser intensity, frame repetition rate, and acquisition time—to achieve a desired level of precision in both fluorophore weight and fluorescence lifetime estimation.

## 4. Discussion

In this study, we developed an uncertainty model for phasor-based time-domain FLIM and quantitatively analyzed the effect of shot noise on the estimation of component weights and fluorescence lifetimes. Our results show that shot noise introduces significant deviations in phasor coordinates, leading to systematic biases in weight and lifetime estimation. Specifically, each component’s contribution corresponds to a two-dimensional Gaussian probability distribution in the phasor plot, which directly impacts the accuracy of component fraction determination. For biological samples exhibiting multiple fluorescence lifetimes, this model enables pixel-wise evaluation of uncertainty in lifetime estimation and its associated component weights, offering a more informed interpretation of FLIM data.

In the Result section, we primarily focused on scenarios involving two or three fluorophore components. In the first-harmonic phasor plot, we established three independent equations: the G and S coordinates of the mixture are expressed as the weighted sum of the corresponding coordinates of the pure components, with the constraint that the weight fractions sum to one. These equations provide a mathematically sufficient framework for unmixing up to three underlying components. However, when the number of components increases to four or more, the intrinsic two-dimensional nature of the first-order phasor plot becomes insufficient to uniquely determine the component weights[27]. To overcome this limitation, higher-order harmonics must be incorporated into the phasor analysis, thereby introducing additional independent equations[28]. Once the component weights are resolved—whether using the first harmonic for simple mixtures or higher harmonics for more complex cases— our uncertainty model can be applied to quantify deviations resulting from shot noise. Given that potential weight values correspond to fixed positions in the first-order phasor plot, the uncertainty analysis can still be efficiently conducted using a single covariance matrix derived from the noise characteristics in one phasor plot.

A fundamental assumption in our model is that each pure fluorophore component exhibits single-exponential fluorescence decay. In cases where components follow multi-exponential decay kinetics, they can be mathematically decomposed into a mixture of single-exponential decays, thereby allowing our model to approximate and quantify noise-induced deviations. However, the reliability of this approach depends on the stability of the decomposition process. Any ambiguity or instability in resolving the underlying single-exponential components may introduce additional errors, potentially compromising the accuracy of the uncertainty analysis.

Furthermore, the current implementation primarily accounts for Poisson-distributed shot noise, yet other sources of detector noise, such as dark noise and readout noise, may also contribute to measurement uncertainty. These additional noise factors may affect both the decomposition stability and the robustness of the uncertainty estimates, underscoring the need to incorporate them in future refinements of the model.

Beyond addressing these limitations, the uncertainty model also holds potential as a diagnostic tool for evaluating FLIM systems[29–31]. By quantifying the spread of phasor coordinates under controlled conditions, it may serve as a means to characterize and compare the noise performance of different FLIM setups. This could help identify system-specific deviations arising from instrumental imperfections or environmental fluctuations, offering insight into factors beyond photon statistics that affect measurement quality.

From an application standpoint, our uncertainty model offers valuable guidance for clustering and segmentation tasks in phasor-based FLIM analysis[10–13]. By quantifying the expected size of the uncertainty region in the phasor plot at a given photon count level, the model supports more informed decision-making when defining cluster boundaries or setting segmentation thresholds, especially in low-photon scenarios where variability is higher.

Another important application of our uncertainty model lies in enabling uncertainty-driven adaptive acquisition in FLIM, which can enhance both data quality and imaging efficiency[32]. By providing pixel-wise uncertainty estimates, the model allows researchers to assess measurement reliability and optimize acquisition parameters such as laser power, exposure time, and sampling rate. Specifically, this framework facilitates an adaptive scanning strategy wherein an initial low-dose, rapid scan identifies regions of high uncertainty. These regions can then be selectively re-imaged at higher precision, while areas with low uncertainty are excluded from further acquisition. This targeted, feedback-based approach holds the potential to improve overall imaging efficiency by minimizing redundant measurements, reducing total light exposure, and mitigating photobleaching and phototoxicity—factors that are especially critical for high-speed, three-dimensional live-cell and tissue imaging applications.

## Acknowledgment

We would like to thank Shimon Weiss, X. Michalet, and Debjit Roy, for their help with extracting data from raw FLIM measurement files.

## Data Availability

Data underlying the results presented in this paper may be obtained from the corresponding author upon reasonable request.

## Supplemental document

See Supplement 1 2 3 for supporting content.

