## Supplementary for "Quantifying Uncertainty in Phasor-Based Time-Domain Fluorescence Lifetime Imaging Microscopy"

**Supplementary 1**

For an ideal FLIM measurement without noise, the phasor coordinates $(G,S)$ at a given pixel are computed by applying a normalized Fourier transform to the fluorescence decay histogram:

$$G=\frac{\sum_{i} N_{box,i}\cos\left( \omega t_{i} \right)}{\sum_{i} N_{box,i}} , (1)$$

$$S=\frac{\sum_{i} N_{box,i}\sin\left( \omega t_{i} \right)}{\sum_{i} N_{box,i}} , (2)$$

where $N_{box,i}$ represents the photon count detected at time bin $t_{i}$, and $\omega$ is the angular frequency corresponding to the analysis temporal histogram range, usually set to the first harmonic. However, due to shot noise, the observed photon count $N_{noisy,i}$ in each time bin follows a Poisson distribution:

$$N_{noisy,i}\sim Poisson\left( N_{box,i} \right) . (3)$$

A defining property of the Poisson distribution is that its variance equals its mean:

$$Var\left[ N_{noisy,i} \right]=N_{box,i} . (4)$$

Shot noise causes phasor coordinates to fluctuate randomly, leading to systematic deviations in lifetime estimation and fluorophore unmixing.

To quantitatively analyze these deviations, $G_{noisy}$ is derived from the histogram of each time bin using Equation (5), with its variance given by Equation (6):

$$G_{noisy}=\frac{\sum_{i} N_{noisy,i}\cos\left( \omega t_{i} \right)}{\sum_{i} N_{noisy,i}} . (5)$$

$$Var\left[ G_{noisy} \right]=\frac{Var\left[ \sum_{i} N_{noisy,i}\cos\left( \omega t_{i} \right) \right]}{\left( \sum_{i} N_{noisy,i} \right)^{2}} . (6)$$

Using error propagation theory, the variance of a function is approximated via a first-order Taylor expansion, which quantifies the sensitivity of output uncertainty to input fluctuations **(Supplementary 2)**. By simplifying the numerator of Equation (6) and substituting Equation (4), the following expression is obtained:

$$\mathrm{Numerator}=\sum_{i} Var\left[ N_{noisy,i} \right]{cos}^{2}\left( \omega t_{i} \right)=\sum_{i} N_{box,i}{cos}^{2}\left( \omega t_{i} \right) . (7)$$

Since $G_{noisy}$ represents a normalized weighted sum, its variance is influenced not only by noise at individual points but also by how each term contributes to deviations from the expected value, represented as $\cos\left( \omega t_{i} \right)-G$. Therefore, the numerator is further refined as:

$$\mathrm{Numerator}= \sum_{i} N_{box,i}{(\cos\left( \omega t_{i} \right)-G)}^{2} . (8)$$

We assume that the total photon count in a pixel remains approximately constant before and after noise is added. Consequently, the denominator of Equation (6) can be expressed as:

$$\mathrm{Denominato}r{=\left( \sum_{i} N_{noisy,i} \right)}^{2}{=\left( \sum_{i} N_{box,i} \right)}^{2} . (9)$$

By combining Equations (8) and (9), expressions for the variance and covariance **(Supplementary 3)** of the normal distributions for $G$ and $S$ under shot noise are derived:

$$Var\left[ G_{noisy} \right]=\frac{\sum_{i} N_{box,i}\left( \cos\left( \omega t_{i} \right)-G \right)^{2}}{\left( \sum_{i} N_{box,i} \right)^{2}} , (10)$$

$$Var\left[ S_{noisy} \right]=\frac{\sum_{i} N_{box,i}\left( \sin\left( \omega t_{i} \right)-S \right)^{2}}{\left( \sum_{i} N_{box,i} \right)^{2}} , (11)$$

$$\mathrm{Cov}\left[ G_{noisy},S_{noisy} \right]=\frac{\sum_{i} N_{box,i}(\cos\left( \omega t_{i} \right)-G)(\sin\left( \omega t_{i} \right)-S)}{\left( \sum_{i} N_{box,i} \right)^{2}} . (12)$$

Thus, each ideal histogram with known pure component weights corresponds to a noise-affected two-dimensional normal distribution, centered along the line connecting the two pure components. For histograms affected by noise, deviations cause points in the phasor plot to belong to multiple two-dimensional normal distributions. The probability of a point belonging to a specific normal distribution $p_{i}$ is given by Equation (13):

$$p_{i}=\frac{1}{2\pi\sqrt{det(\boldsymbol{C}_{i})}}\exp\left( -\frac{1}{2}\left( \boldsymbol{x-}\boldsymbol{\mu}_{\boldsymbol{i}} \right)^{T}\boldsymbol{C}_{i}^{-1}\left( \boldsymbol{x-}\boldsymbol{\mu}_{\boldsymbol{i}} \right) \right) . (13)$$

Here, $\boldsymbol{\mu}$ denotes the mean $(G, S)$ values, $\boldsymbol{C}$ is the covariance matrix of the normal distribution, and $\boldsymbol{x}$ represents the measured noisy data in the phasor plot. By normalizing these probabilities across all distributions, the weight probability for each pure component can be expressed as:

$$P_{i}=\frac{p_{i}}{\sum_{j=1}^{n} p_{j}} . (14)$$

**Supplementary 2**

Assume that $y=f(x_{1},x_{2},\ldots,x_{n})$ is a function of the input variables $x_{1},x_{2},\ldots,x_{n}$, each with an associated uncertainty, expressed as $Var[x_{i}]$. The function can be expanded using a first-order Taylor series at the point $(x_{1},x_{2},\ldots,x_{n})$:

$$f\left( x_{1}+\Delta x_{1},x_{2}+\Delta x_{2},\ldots x_{n}+\Delta x_{n} \right)\approx f\left( x_{1},x_{2},\ldots,x_{n} \right)+\sum_{i=1}^{n} \frac{\partial f}{\partial x_{i}}\Delta x_{i} . (15)$$

Where $\Delta x_{i}$ represents small changes in the input $x_{i}$, and $\frac{\partial f}{\partial x_{i}}$ denotes the partial derivative of $f$ with respect to $x_{i}$. Therefore, the change in output $\Delta y$ can be approximated as:

$$\Delta y=\sum_{i=1}^{n} \frac{\partial f}{\partial x_{i}}\Delta x_{i} . (16)$$

Due to the definition and translation invariance property of variance, $Var\left[ y \right]$ can be expressed as:

$$Var\left[ y \right]=Var\left[ \Delta y \right]=E\left[ \left( \Delta y \right)^{2} \right]-\left( E\left[ \Delta y \right] \right)^{2} . (17)$$

Since the first-order Taylor expansion ignores higher-order terms, it can be assumed that $E[\Delta y]\approx0$. Allowing Equation (17) to be simplified as:

$$Var\left[ y \right]=E\left[ \left( \Delta y \right)^{2} \right] . (18)$$

By substituting Equation (16) into Equation (18):

$$Var\left[ y \right]=E\left[ \left( \sum_{i=1}^{n} \frac{\partial f}{\partial x_{i}}\Delta x_{i} \right)^{2} \right]=E\left[ \sum_{i=1}^{n} \left( \frac{\partial f}{\partial x_{i}} \right)^{2}\left( \Delta x_{i} \right)^{2}+\sum_{i\neq j} \frac{\partial f}{\partial x_{i}}\frac{\partial f}{\partial x_{j}}\Delta x_{i}\Delta x_{j} \right] . (19)$$

If the inputs $x_{1},x_{2},\ldots,x_{n}$ are independent, then for $i\neq j$, $E\left[ \Delta x_{i}\Delta x_{j} \right]=0$. Given the uncertainty in $x_{i}$, $Var\left[ x_{i} \right]=E[{(\Delta x_{i})}^{2}]$. The variance of output $y$ can be finally expressed as:

$$Var\left[ y \right]=E\left[ \sum_{i=1}^{n} \left( \frac{\partial f}{\partial x_{i}} \right)^{2}\left( \Delta x_{i} \right)^{2} \right]=\sum_{i=1}^{n} \left( \frac{\partial f}{\partial x_{i}} \right)^{2}Var[x_{i}] . (20)$$

**Supplementary 3**

From the definition of covariance,

$$\mathrm{Cov}\left[ G_{noisy},S_{noisy} \right]=E\left[ (G_{noisy}-G)(S_{noisy}-S) \right] . (21)$$

Where:

$$G_{noisy}-G=\frac{\sum_{i} {(N}_{noisy,i}-N_{box,i})\cos\left( \omega t_{i} \right)}{\sum_{i} N_{noisy,i}} , (22)$$

$$S_{noisy}-S=\frac{\sum_{i} {(N}_{noisy,i}-N_{box,i})\sin\left( \omega t_{i} \right)}{\sum_{i} N_{noisy,i}} , (23)$$

Therefore:

$$E\left[ (G_{noisy}-G)(S_{noisy}-S) \right]=E\left[ \left( \frac{\sum_{i} {(N}_{noisy,i}-N_{box,i})\cos\left( \omega t_{i} \right)}{\sum_{i} N_{noisy,i}} \right)\left( \frac{\sum_{j} {(N}_{noisy,j}-N_{box,j})\sin\left( \omega t_{j} \right)}{\sum_{j} N_{noisy,j}} \right) \right] . (24)$$

Expanding the numerator:

$$E\left[ {(N}_{noisy,i}-N_{box,i}){(N}_{noisy,j}-N_{box,j}) \right]=E\left[ N_{noisy,i}N_{noisy,j} \right]-N_{box,i}E\left[ N_{noisy,j} \right]-N_{box,j}E\left[ N_{noisy,i} \right]+N_{box,i}N_{box,j}=E\left[ N_{noisy,i}N_{noisy,j} \right]-N_{box,i}N_{box,j} . (25)$$

For the case where $i=j$:

$$E\left[ {N_{noisy,i}}^{2} \right]=Var\left[ N_{noisy,i} \right]+\left( E\left[ N_{noisy,i} \right] \right)^{2}=N_{box,i}+{N_{box,i}}^{2} , (26)$$

$$E\left[ {(N}_{noisy,i}-N_{box,i}){(N}_{noisy,j}-N_{box,j}) \right]=N_{box,i} . (27)$$

when $i\neq j$,

$$E\left[ N_{noisy,i}N_{noisy,j} \right]=E\left[ N_{noisy,i} \right]\cdot E\left[ N_{noisy,j} \right]=N_{box,i}N_{box,j} , (28)$$

$$E\left[ {(N}_{noisy,i}-N_{box,i}){(N}_{noisy,j}-N_{box,j}) \right]=0 . (29)$$

Therefore,

$$\mathrm{Cov}\left[ G_{noisy},S_{noisy} \right]=\frac{\sum_{i} N_{box,i}\cos\left( \omega t_{i} \right)\sin\left( \omega t_{i} \right)}{\left( \sum_{i} N_{box,i} \right)^{2}} . (30)$$

Similarly, since covariance depends not only on Poisson noise but also on the weight distribution at each time point, it is further refined as:

$$\mathrm{cov}\left[ G_{noisy},S_{noisy} \right]=\frac{\sum_{i} N_{box,i}(\cos\left( \omega t_{i} \right)-G)(\sin\left( \omega t_{i} \right)-S)}{\left( \sum_{i} N_{box,i} \right)^{2}} . (31)$$
